# Empowering high-throughput high-content analysis of 3D tumor models: Open-source software for automated non-confocal image analysis

**DOI:** 10.1101/2024.02.20.581231

**Authors:** Noah Wiggin, Carson Cook, Mitchell Black, Ines Cadena, Salam Rahal-Arabi, Kaitlin Fogg

**Affiliations:** School of Electrical Engineering and Computer Science, Oregon State University, Corvallis, OR; School of Chemical, Biological, and Environmental Engineering, Oregon State University, Corvallis, OR

**Keywords:** bioinformatics, computational biology, tumor microenvironment, pharmacokinetics, tissue engineering, hydrogel

## Abstract

**Purpose:** The primary aim of this study was to develop an open-source Python-based software for the automated analysis of dynamic cell behaviors in three-dimensional tumor models using non-confocal microscopy. This research seeks to address the existing gap in accessible tools for high-throughput analysis of cancer and endothelial cell dynamics *in vitro*, facilitating the rapid assessment of drug sensitivity.

**Methods:** Our approach involved annotating over 1000 2 mm Z-stacks of cancer and endothelial cell co-culture model and training machine learning models to automatically calculate cell coverage, cancer invasion depth, and microvessel dynamics. Specifically, cell coverage area was computed using focus stacking and Gaussian mixture models to generate thresholded Z-projections. Cancer invasion depth was determined using a deep neural network binary classification model, measuring the distance between Z-planes with invaded cells. Lastly, microvessel dynamics were assessed through a U-Net Xception-style deep learning model for segmentation, a disperse algorithm for network graph representation, then persistent homology to quantify microvessel length and connectivity. Finally, we reanalyzed an image set from a high-throughput drug screen involving a chemotherapy agent on a 3D cervical and endothelial co-culture model.

**Results:** The software accurately measured cell coverage, cancer invasion, and microvessel length, yielding drug sensitivity IC_50_ values with a 95% confidence level compared to manual calculations. Additionally, it significantly reduced the image processing time from weeks down to hours.

**Conclusions:** Our free and open source software offers an automated solution for quantifying 3D cell behavior in tumor models using non-confocal microscopy, providing the broader Cellular and Molecular Bioengineering community with an alternative to standard confocal microscopy paired with proprietary software.

## 1. Introduction

Cancer is a complex disease characterized by uncontrolled cell growth, invasion, and metastasis (Bielenberg & Zetter, 2015). Understanding the mechanisms underlying these processes is critical for developing effective treatment strategies. 3D tumor models are a useful tool to capture the interactions between tumor cells and the surrounding microenvironment, which play a key role in tumor progression (Bray & Werner, 2018). However, there remains a lack of open-source image analysis software for automated assessment of dynamic tumor behaviors like growth, invasion, and angiogenesis (Booij et al., 2019). This is a bottleneck for high throughput imaging experiments, hindering our ability to fully exploit the potential of 3D models to study these cell behaviors under various experimental conditions.

Common bioimage analysis software tools such as Fiji ImageJ (Schindelin et al., 2012) provide a diverse array of tools and a graphical user interface for detailed analysis of images. However, the manual work required to use these tools increases in proportion to the number of images in the dataset, limiting their application for high throughput image analysis. Recent advancements in machine learning (ML) for computer vision have created new pathways for automated and accurate analysis of biological images. Deep learning has shown exceptional promise in image recognition and segmentation tasks in the domain of biomedical image analysis. These techniques have been used successfully in various applications, such as in the classification of invasion depth in esophageal squamous cell carcinoma (Nakagawa et al., 2019) and in cell coverage area analysis in time-lapse fluorescence microscopy (Rundo et al., 2020).

Deep learning models such as U-Net (Chen et al., 2020; Ronneberger et al., 2015) have demonstrated high accuracy in vessel segmentation tasks. Further accuracy may be obtained by considering the structural characteristics of a vessel network, such as graphical connectivity (Shin et al., 2019). Additionally, analyzing the structural properties of microvascular networks reveals phenotypic information of significant biological relevance (Corliss et al., 2019). An emerging area in the quantitative analysis of biological networks is the application of topological data analysis techniques (Dey & Wang, 2022; Edelsbrunner & Harer, 2010). In biological imaging, topological data analysis tools such as persistent homology have been used to characterize and quantify complex structures, such as the branching architecture of microvessel networks in brain artery trees (Bendich et al., 2016) and neuronal structures (Kanari et al., 2018). In recent years, topological data analysis has been applied to characterize the structural and functional characteristics of vessel networks, including tumor vascular networks (Nardini et al., 2021; Stolz et al., 2022).

Despite these advancements, there remains a need for comprehensive, open-source tools designed explicitly for measuring cell behaviors of 3D models in high throughput non-confocal imaging experiments. The development of such tools would facilitate studies into cell-microenvironment interactions in 3D cultures and contribute towards understanding the effect of the tumor microenvironment on cancer progression.

Here we introduce an open-source software package for the high-throughput analysis of cancer and endothelial cell dynamics in hydrogels. We applied this software to 3D multilayer multicellular models of cervical and endometrial cancer cells co-cultured with human microvascular endothelial cells (hMVEC). Our software package automates the quantification of cell coverage area, invasion depth, and microvessel formation, enabling rapid and accurate assessment of phenotypic cell responses in 3D tumor models.

## 2. Materials and Methods

### 2.1. Cell lines and reagents

Unless stated, all reagents were purchased from ThermoFisher (Waltham, MA). Human microvascular endothelial cells (hMVEC) were purchased from Lonza (hMVEC 33226, Walkersville, MD) and used without additional characterization. Cells were expanded in EGM-2 MV media (EBM-2 supplemented with Lonza’s SingleQuot supplements: hydrocortisone, human basic fibroblast growth factor (FGF2), human vascular endothelial growth factor (VEGF), human insulin-like growth factor (IGF), human epidermal growth factor (EGF), ascorbic acid, and gentamycin) and further supplemented with 5% fetal bovine serum (FBS) until used at passage 5. Human cervical cancer cell lines SiHa (ATCC® HTB-35™) and Ca Ski (ATCC CRM-CRL-1550) and human endometrial cancer cell line HEC-1A (ATCC HTB-112™) were purchased from ATCC (Manassas, VA) and used without additional characterization. The cells were cultured in 1% penicillin–streptomycin (Sigma-Aldrich, St. Louis, MO, USA) and 10% fetal bovine serum maintaining Eagle’s Minimum Essential Medium (EMEM, ATCC), RPMI-1640 Medium (ATCC), and McCoy’s 5A medium (ATCC) respectively until used at passage 5. All cell types were expanded in standard cell culture conditions (37℃, 21% O2, 5% CO2) and subcultured before they reached 80% confluency.

### 2.2. Multilayer Hydrogel Fabrication

Multilayer multicellular constructs for cervical cancer cells and endometrial cancer cells were prepared following the same methodology as previously reported (Cadena et al., 2023). Constructs were fabricated in specialized µ-Plate Angiogenesis 96-wells, ibidi Treat (ibidi, Munich, Germany) by layering 10 µL of the bottom hydrogel formulation into each well, and 20,000 CellTracker Green labeled (CellTracker™ Green CMFDA Dye) hMVEC cells in 40 µL of EGM-2 MV were pipetted on top of each gel. The endothelial cells were allowed to attach for four hours at 37°C and 5% CO_2_. The media was then carefully removed, and 25 µL of the top hydrogel formulation was pipetted on top. Finally, cell tracker Red-labeled cancer cells (CellTracker™ Red CMTPX Dye) were seeded on top of each gel at 10,000 cells in 25 µL of media. As a control, multilayer Matrigel constructs were made using the same methods described above, except the custom hydrogel formulations were replaced with growth factor deficient (GFD) Matrigel basement membrane matrix (9.2 mg/mL protein concentration, Corning, MA, USA) in both layers. The media was carefully changed every 12 hours, and the plates were incubated at 37°C and 5% CO_2_ in a BioSpa live cell analysis system (Agilent Technologies, Santa Clara, CA) for 45 hours.

### 2.3. Phenotypic cell response to Paclitaxel

Using the optimized cervical cancer construct (Cadena et al., 2023), we measured the phenotypic cell responses: microvessel length, cervical cancer invasion, endothelial cell coverage, and cancer cell coverage in endothelial cells (hMVEC), and the cervical cancer cells (SiHa and Ca Ski) cultured the multilayer multicellular models and treated with the chemotherapy agent Paclitaxel for 48 hours. Paclitaxel was a kind gift from the Oregon State University College of Pharmacy High-Throughput Screening Services Laboratory. Cells were imaged every 12 hours, and the cell response to the drug was reported at 24 hours of treatment. The dose-response analysis of Paclitaxel ranged from 0.008 to 25 μM, and dimethyl sulfoxide (DMSO) was used as a vehicle. Paclitaxel and DMSO were dispensed in the wells using an automated liquid handler (D300e Digital Dispenser, Hewlett Packard, Corvallis, OR). The dose response-inhibition curves were calculated using Prism 8.2.1 software (GraphPad, San Diego, CA). We then evaluated the IC_50_ values in terms of phenotypic responses. While IC_50_ values are classically defined as the concentration that inhibits 50% of the cells from proliferating or the concentration that kills 50% of the cells, here we defined the IC_50_ values as the concentration of drug that reduced either cancer invasion, cancer coverage, microvessel length, or endothelial cell coverage to 50% of the value observed in the absence of drug. Lastly, we compared previously reported IC_50_ values using Fiji ImageJ (NIH, Bethesda, MD) and the Gen5 software (Agilent Technologies) (Cadena et al., 2023) with the described image analysis tools.

### 2.4. Image acquisition in multilayer multicellular model

We have previously demonstrated that our multilayer multicellular 3D *in vitro* model can support the phenotypic cell response over time (Cadena et al., 2023). The phenotypic responses were defined as cancer cell coverage area, cancer cell invasion, endothelial cell coverage area, and endothelial microvessel length. Two-channel 85 µm z-stack images were taken using a Cytation 5 V3.14 cell imaging multi-mode reader (Agilent Technologies). A total of 21 z-stacks per well and per experiment were recorded and used for post-processing to calculate the cell phenotypic response in each hydrogel model.

### 2.5. Software package for automated image analysis

We developed a Python software package that contains four automated image analysis tools designed to facilitate the quantification of phenotypic cell response in a high-throughput setting. Our Python software package contains automated image analysis tools for assessing microvessel formation, cell coverage, and cancer invasion depth. This open-source software package can be used as either a standalone command-line utility or as an integrated component within other software programs.

Comprehensive documentation detailing installation and usage instructions can be found in our GitHub repository: https://github.com/fogg-lab/tissue-model-analysis-tools. Additionally, we have provided an interactive demonstration of each analysis tool in the supplementary materials, which can be accessed via the Colab notebook link: https://colab.research.google.com/github/fogg-lab/tissue-model-analysis-tools/blob/main/notebooks/analysis_demo.ipynb.

### 2.6. Making Z-projection images

For manual calculations, the Cytation 5 a Z-projection function was used to combine the 21 Z-stacks into a 2D Z-projection. Using a focus stacking with a maximum filter size of 11 px, we captured the cell response in one 2D image. For automated calculations, we employed five established projection methods to compute a Z-projection of input Z-stacks. These methods include focus stacking, minimum pixel intensity, maximum pixel intensity, median pixel intensity, and average pixel intensity. Users can select the most suitable method for computing Z-projections of their samples, depending on the characteristics of their dataset. To compare the phenotypic cell response between manual inspection and automated analysis, we chose the focus stacking approach, generating an output comparable to that of Gen5.

### 2.7. Manual inspection of cell coverage area

After computing the Z-projections with Gen5, images were processed with NIH Fiji-ImageJ (NIH, Bethesda, MD). Cell coverage was measured for each cell type at every time point by calculating the area within the well covered by cells and dividing it by the total well area. A macro recorded code was generated to process a batch of images over time, code is described in **Supplemental Fig 1**. Cell coverage was reported as a percentage of the endothelial cells in the well as well as the percentage of cancer cells in the well.

### 2.8. Automated computation of cell coverage area

In the software package, we implemented a procedure that incorporates a two-component Gaussian Mixture Model (GMM). Two Gaussian curves are fit to the pixel intensities in the image, with one curve fit to the foreground pixels and the other fit the background pixels. The foreground pixels are defined as the bright regions in the image (high pixel intensities), indicating cells, while the background comprises the darker regions without cells (low pixel intensities).

After fitting the GMM, we then compute cell area by thresholding the image based on a cutoff pixel intensity calculated as *μ_foreground_* + *λ* × *σ_foreground_*. Here, *μ_foreground_* and *σ_foreground_* represent the estimated mean and standard deviation of the foreground intensity, which are parameters extracted from the GMM’s foreground component. *λ* is a tunable parameter used as a multiplier of the foreground standard deviation. When *λ* = 0, pixels with intensities greater than *μ_foreground_* pass the threshold. When *λ* > 0, fewer pixels pass the threshold. When *λ* < 0, more pixels pass the threshold. This approach is effective for images containing a bimodal distribution of pixel intensities, regardless of the overall brightness and contrast of each image. Similar to manual inspection, cell coverage was then calculated as the cell area divided by the total area of the well. (**Figure 1**). The well boundary was determined by estimating the parameters of a superellipse that fits around the Canny edges of the image. If this failed due to background noise near the image border, or due to a lack of detected Canny edges, we used a circular boundary centered on the image.

**Figure 1.**
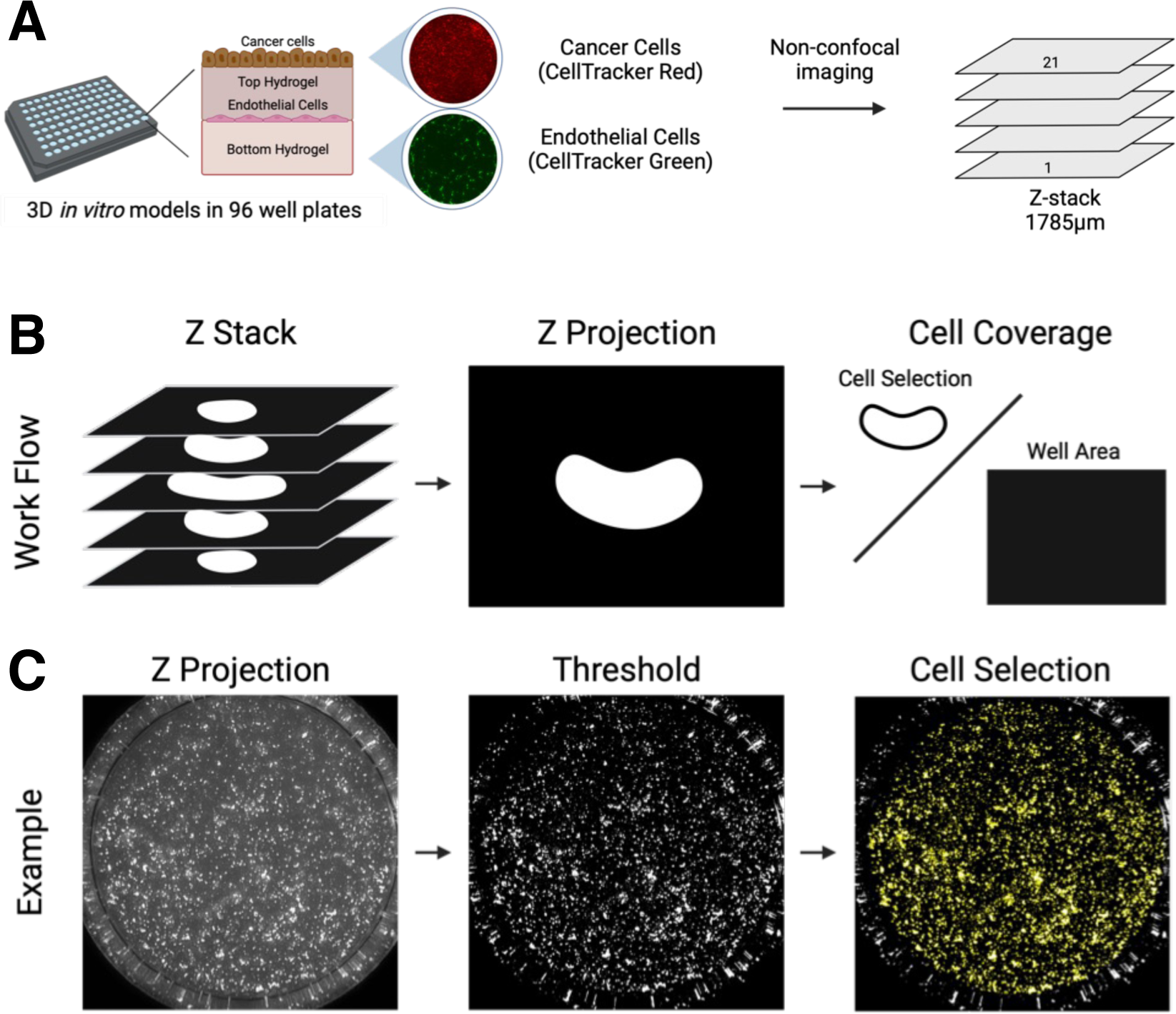
Schematic representation of image acquisition and cell thresholding. (**A**) CellTracker Green labeled human microvascular endothelial cells (hMVEC) were cocultured in different 3D *in vitro* hydrogel models with CellTracker Red labeled cancer cells (SiHa, CaSki, or HEC-1A). Constructs were imaged every three hours for 48 hours with two different channels (GFP and Texas Red) and a 1785 µm Z-stack was taken for each well to capture full construct height. (**B**) Schematic demonstrating Z-projection followed by cell coverage calculation (C) Example of how Z-projection images are used to calculate cell coverage.

### 2.9. Manual inspection of cell invasion depth

Cell invasion depth was defined as the deepest point of cancer cell invasion after 24 hours, and 48 hours of cultured in the 3D multilayer model. This was achieved by annotating each Z-stack image set and calculating the difference between cell invasion from time 0h, 24h and 48h. Using Gen5, we manually inspected each well and z-stack to track cell movement over time.

### 2.10. Automated computation of cell invasion depth

We use a binary classifier deep neural network based on the ResNet50 architecture to estimate the depth of invasion in a given Z-stack. We trained the classifier on a dataset of 997 images, split into two classes - *invasion* and *no invasion*. 848 of these images were invaded, and 149 were not invaded (**Figure 2**). During training, we augmented the dataset by applying random flips and rotations to the samples. The binary classifier assesses each image in a Z-stack to determine whether it displays a sufficient amount of in-focus cell area to be considered as demonstrating invasion. A Z-stack of input images *Z* = (*z_k_*, *z*_*k*-1+_, … , *z*_0_) is processed by the invasion depth analysis system, which outputs two results:

1. A collection of probabilities, *p̂*, showing the model’s confidence that invasion has been identified:

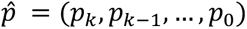
2. A collection of classifications, *ŷ_p_* thresholded at a given value (typically *p_i_* > 0.5):

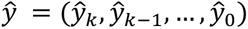

**Figure 2:**
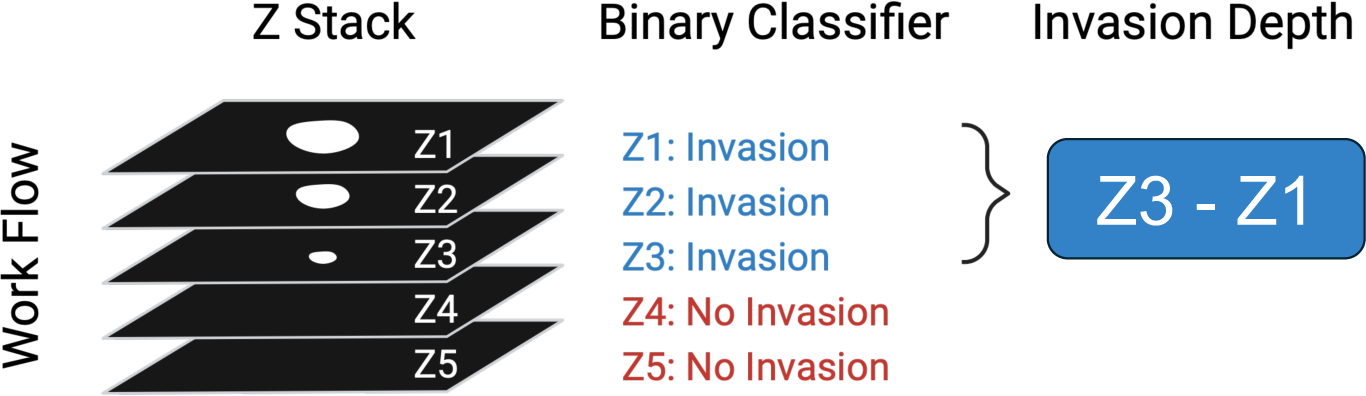
Schematic representation of invasion probability determination. For each input image, a binary classifier calculates the probability of invasion, with a probability greater than or equal to 0.5 indicating that the sample is classified as “Invaded.” Invasion depth is then calculated as the distance between the top z plane and the deepest Z-plane with invasion.

### 2.11. Manual quantification of microvessel formation

Endothelial microvessel length was quantified by measuring the average length of the microvessels found in a well at each time point using NIH Fiji-ImageJ. We use a macro code to determine the number of microvessels, total branch length and average length of each branch in each set of Z-projected images (**Supplemental Fig 2**). The average branch length was reported in micrometers.

### 2.12. Semantic binary segmentation

We trained a U-Net Xception-style model (Chollet, 2016; Ronneberger et al., 2015) to segment the microvessels from the background of an image. The target output from this process is a probabilistic (non-thresholded) segmentation of the image, where each predicted value is the probability that the pixel at that position is part of a vessel. To train, validate, and test the segmentation model, we prepared a dataset of fifty manually annotated images. Each annotated image in the dataset consisted of the original image and its corresponding annotation in the form of a binary segmentation mask. We produced annotations for fifty images using an interactive segmentation program (Fogg Lab, 2022) based on the RITM: Interactive Segmentation codebase (Sofiiuk et al., 2021). We split the annotated images into three datasets - a training set of thirty samples, a validation set of ten samples, and a test set of ten samples.

At training time, we applied a series of image transformations to each training sample as a form of data augmentation. These transformations improved the model’s convergence and were necessary for the model to generalize to images outside the training dataset. The transformations and the probabilities for applying each transformation were chosen experimentally. We applied the following image transformations to the training samples: rotate at a random angle (*p* = 0.5), crop to a random 512×512 patch (*p* = 1.0), flip horizontally (*p* = 0.25) or vertically (*p* = 0.25), alter brightness and contrast (*p* = 0.7), apply either multiplicative noise (*p* = 0.4) or gaussian blur with noise (Buslaev et al., 2020) (*p* = 0.4), and apply elastic deformations (Bloice et al., 2017) (*p* = 0.85). The transformed samples were subsequently resampled to the target size of 320×320. Lastly, each sample was normalized with the mean and standard deviation of the training set. To find suitable hyperparameters to train the segmentation model, we conducted a grid search over a range of options for the filter counts and the initial learning rate. The search space was determined experimentally, consisting of seven options for the initial learning rate and three options for the filter counts. For our dataset, the best learning rate was 0.001, and the best filter counts were (64, 128, 256, 512).

We built a model with the best filter counts and initial learning rate trained it for fifty epochs. Each epoch consisted of ⌊1500/*batchsize*⌋ training steps, and ⌊500/*batchsize*⌋ validation steps. At each training step, *batchsize* samples were chosen at random from the training dataset. At each validation step, *batchsize* samples were chosen at random from the validation data2set. To obtain probabilistic segmentations of full resolution images using the model, which takes 320×320 image patches as input, images were divided into overlapping 512 × 512 patches. Each patch was downsampled to the target size, normalized, and processed by the model, resulting in a probabilistic segmentation of each patch. Subsequently, the overlapping tiled predictions were blended together (Vooban Inc., 2017). The blended prediction value for each overlapping pixel position was computed as the average value of the corresponding pixel in the overlapping tiles.

### 2.13. Post-processing raw segmentations prior to graph extraction

We post-processed the probabilistic segmentations to remove background noise and enhance vessel centerlines. First, we removed circular regions and small disconnected components from the segmentation. To find these components, we first computed a binary segmentation mask by thresholding the probabilistic segmentation at *p* = 0.5, and assigned a component ID to the pixels within each connected component of the mask. We calculated circularity of each component using the formula, 4*π* × *area* ÷ *perimeter*^2^, and removed components whose circularity is greater than 0.8. Additionally, we computed a 1-pixel wide skeleton (Zhang, Tongjie Y & Suen, Ching Y., 1984) of each component of the segmentation mask, and removed components whose skeleton contained no junctions, which indicates that there are no attached branches. After removing these spurious components, we applied the filtered segmentation mask to the probabilistic segmentation to remove the background noise. Secondly, we applied another post-processing operation to the probabilistic segmentation to increase the values of the pixels on the expected centerlines of the vessels, relative to the pixel intensities on the edges of the vessels. This step is important because our subsequent processing methods assume the bright ridges on the image represent the centerlines of vessels. This is not the case for the raw probabilistic segmentation, as the class probabilities tend to be uniform along a cross-section of a vessel. To enhance the centerlines of the predicted vessel regions, we computed the medial axis of the segmentation mask and multiplied the segmentation class probabilities by weights computed by a custom distance function. The weight for each pixel was calculated as a function of its Euclidean distance from the nearest point on the medial axis pixel and the Euclidean distance to the nearest background point, per the formula, 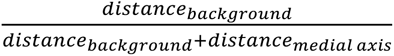. This function effectively computes a distance transform relative to the local vessel width. The probabilistic segmentation was multiplied by these weights to bring vessel centerlines into focus, isolating the appropriate signal for subsequent graph extraction.

### 2.14. Graph Extraction

To calculate statistics of the endothelial cell microvessel formation, we needed a way of extracting a graph representation (i.e. a list of vertices and edges) from the image. We used the Discrete Persistent Structures Extractor (DisPerSE) algorithm (Sousbie, 2011) to extract a graph representation of a microvessel network. The DisPerSE algorithm was originally developed as a way of extracting a representation of the “cosmic web’’ of galaxies. It has since been used in Computational Biology as a way of extracting biological networks from image data. For example, it has been used to extract representation of neuronal data from images of neurons. As the DisPerSE algorithm is heavily inspired by the mathematical field of Morse Theory, we refer to the graph representation computed by the DisPerSE algorithm as the *Morse skeleton*.

Intuitively, the DisPerSE algorithm works as follows. We can think of a grayscale image as a “mountain range” in three dimensions, where the x and y coordinates of a pixel are its x and y coordinates in an image, and the z coordinate of a pixel is its brightness. The graph returned by Disperse are the “mountain ridges” connecting different peaks in the mountain range. However, the mountain range of an image may have many more ridges than the network depicted in the image, as noise in the image can add many different peaks of similar height around a “true” peak. Thus, Disperse performs a smoothing step to remove multiple peaks of similar height by “canceling” two nearby peaks and the saddle connecting them.

The key challenge of extracting a graph from our dataset was that the distribution of endothelial cells are non-uniform; rather than forming a line, microvessels are formed and changed over time and based on the material that the endothelial cells are seeded. Therefore, images of microvessels are not of uniform brightness. The DisPerSE algorithm is an appropriate method for extracting a representation of a network from an image as it is well-suited for non-uniform data, as the DisPerSE algorithm can “connect the dots” between bright regions of the image separated by dimmer regions.

However, this ability to “connect the dots” can also be a disadvantage, as DisPerSE will try to connect any bright region to the microvessel network. This can be problematic as images often contain cells which are not a part of the main microvessel network. In this case, DisPerSE will add a path connecting these isolated cells to the main microvessel network. To prevent DisPerSE from returning an overly connected network, we filtered for background noise as described in the previous section.

We then simplified the network extracted by DisPerSE in two ways. First, we removed branches shorter than 10 µm, as these branches are often the result of noise in the original image or segmentation mask. Furthermore, we found that this step was key for returning a network that more closely agreed with human reviewers in terms of number of branches and average microvessel length. Second, we performed moving average smoothing on each of the branches. This is because the graph extracted by the DisPerSE algorithm was a subgraph of the grid connecting neighboring pixels. This means that all of its edges were either vertical, horizontal, or at a 45 degree angle. We found that smoothing the branches gave a graph representation that more accurately traces the underlying microvessel network.

### 2.15. Topological Data Analysis

Once we had a graph representation of a microvessel network, our next step was to characterize the branching structure of the microvessels. For example, we wanted to be able to distinguish between networks by the number and lengths of branches. For this task, we needed a way of uniquely decomposing a network into a set of branches (**Figure 3**). For this, we relied on the theory of *persistent homology*.

**Figure 3:**
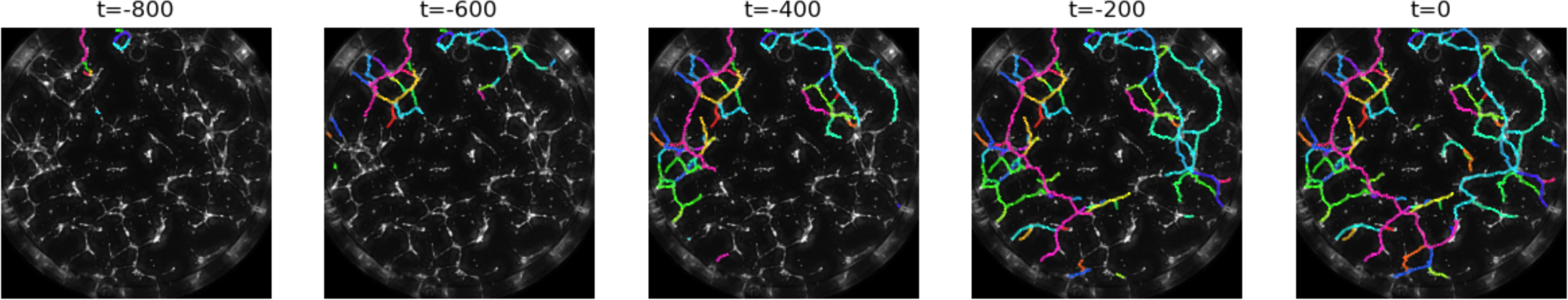
Example of filtration on a Morse skeleton. As the value of t increases, the tree grows from the leaves toward the root. Whenever a new leaf is added, a bar is added to the persistence diagram. When two branches merge, the shorter bar ends in the persistence diagram.

We now present an informal overview of persistent homology. We recommend the book by Edelsbrunner and Harer (Edelsbrunner & Harer, 2010) for a formal introduction to persistent homology. In short, persistent homology is a way of summarizing the topological features of a space across different scales. Generically, persistent homology takes as input a space *X* and a real-valued function *f*: *X* → ℝ. It then considers the ***filtration*** of subspaces *X*_t_ = {*x* ∈ *X* ∶ *f*(*x*) ≤ *t*} as we increase the value *t*. Persistent homology tracks how certain topological properties (such as the number of connected components or loops) changes as we increase the parameter *t*.

The changes in topology are summarized in a ***barcode***. A barcode is a collection of intervals {[*b_i_*, *d_i_*] ∶ 1 ≤ *i* ≤ *m*}. Each ***bar*** [*b_i_*, *d_i_*] describes the lifetime of a specific topological feature. For example, the feature could be a connected component. The value *b_i_* is the real number of the space 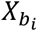 where the feature first appeared, or was ***born***, and the value *d_i_* is the real number of the space 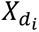 where the feature merged with an older feature, or ***died***. The value (*d_i_* − *b_i_*) is the ***persistence*** of a feature.

In our example, the space *X* is the endothelial network, and the value *f*(*x*) for a point *x* in our network is the negative distance from *x* to a fixed point *r*. It is helpful to visualize the filtration of *f* in the case that our network is a tree and *r* is a root. As we increase the value of *t*, we can imagine the space *X*_t_ growing from the leaves towards the root (**Figure 3**). Each bar [*b_i_*, *d_i_*] corresponds to a branch in a tree. The value *b_i_* is the negative distance of the leaf from the root, and the value *d_i_* is the negative distance from the root at the point where the branch merges with a longer branch. The persistence (*d_i_* − *b_i_*) is the length of the branch. This filtration has previously been applied to the task of quantifying the branching structures of neurons (Kanari et al., 2018).

However, it was not always the case that our microvessel network was a tree. In this case, the barcode of the filtration *f* does not have the same interpretation of decomposing the network into different branches. Specifically, if the network contained a loop, the filtration would not necessarily decompose this loop into two branches. To remedy this, we computed the persistent homology of the shortest path tree from a specified root *r*, where *r* is chosen to be the center of the network. We then used the barcode to provide a summary of the branching structure of the network, reporting the number of branches, total length of the network, and average microvessel length (**Figure 4**). We compared these metrics with the manual measurements outlined in section above. focusing on the measurement of average branch length. Specifically, we evaluated the accuracy of the model predictions by calculating the residuals (i.e. ***observed*** – ***predicted***) and the coefficient of determination (R^2^) for the comparison of manual and automated measurements.

**Figure 4.**
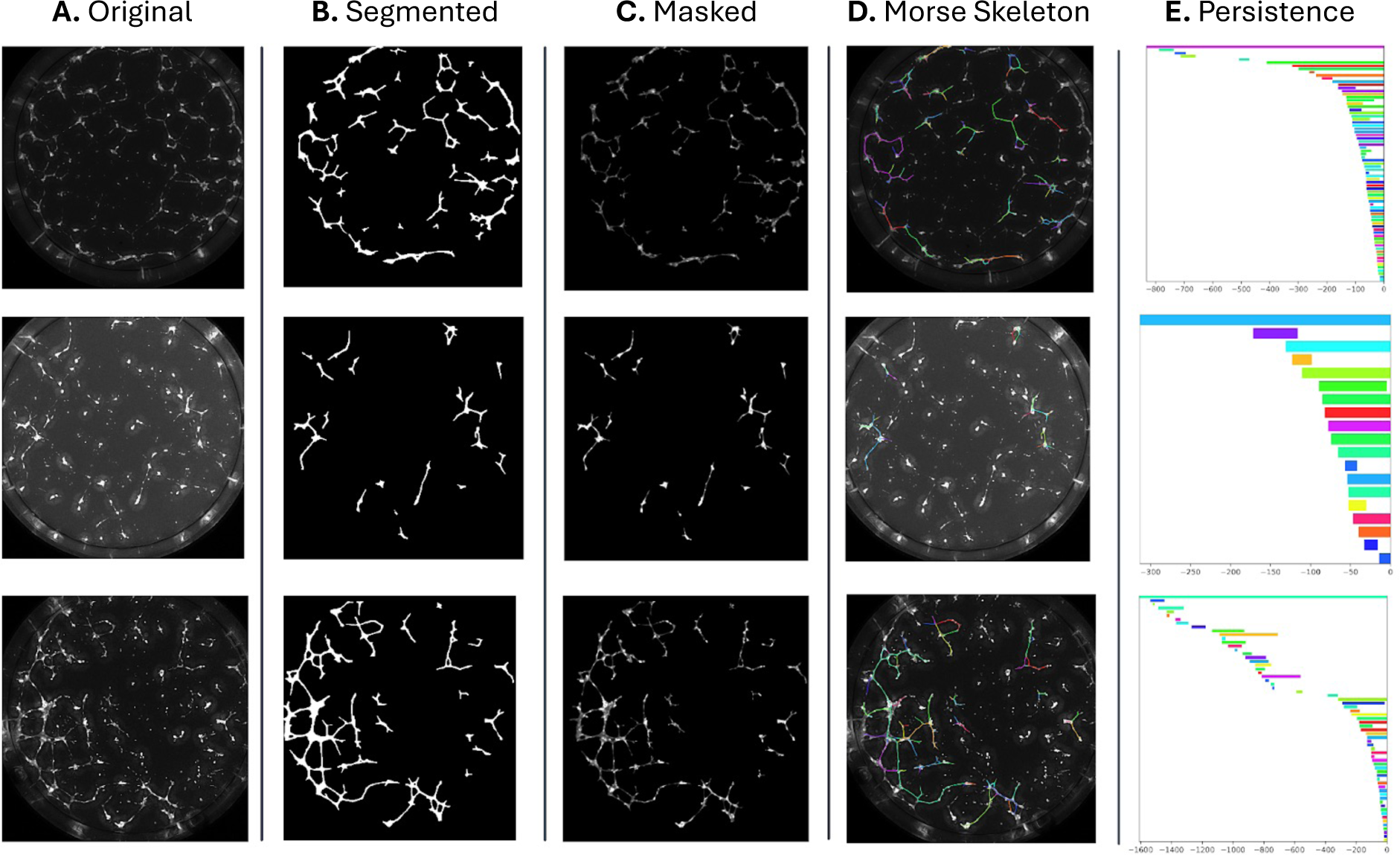
Schematic representation of microvessel network analysis. Input images (**A**) are first processed by a binary segmentation model, which predicts the location of vessels within the image. The raw, probabilistic segmentation (**B**) is then refined by removing blobs and disconnected branches, and the expected vessel centerlines are enhanced by applying multiplicative weights based on a distance transform of the segmentation mask relative to the local vessel width (**C**). The Disperse algorithm is then used to extract a graph representation of the microvessel network (**D**), and a persistence barcode (**E**) is calculated to serve as a topological descriptor of the microvessel network.

## 3. Results

### 3.1. Cell coverage area

To evaluate the accuracy of our software package in estimating cell coverage, we compared the computed area measurements with manual area measurements for the multilayer multicellular models of cervical and endometrial cancers. Randomly selected images ranging from the minimum to the maximum of cell coverage representing all three coculture models were used to compare cell coverage area measured by hand and with our code (**Figure 5**). The coefficient of determination (R^2^) was above 90% for both endothelial cells (**Figure 5A**) and cancer cells (**Figure 5B**).

**Figure 5.**
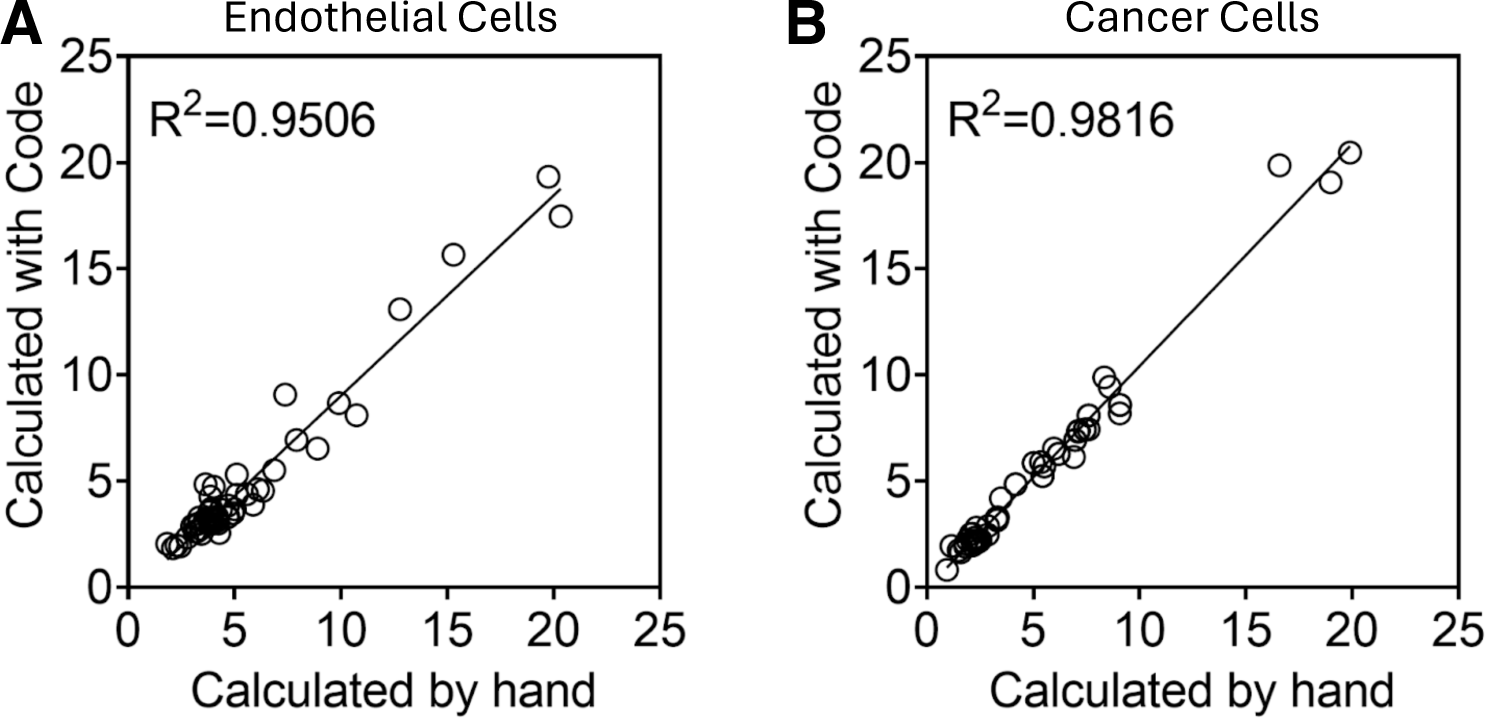
Comparison of Manual measurements and developed code for the percentage of cell coverage. Randomly selected images representing all three coculture models were selected and cell coverage calculated with the code was directly compared to cell coverage calculated by hand. This was performed for (**A**) endothelial cell coverage and (**B**) cancer cell coverage. N=58 images.

### 3.2. Invasion depth

To assess the accuracy of our software package’s invasion depth analysis system, we compared the output of the binary classifier with manual classifications (**Figure 6**). Randomly selected images ranging from the minimum to the maximum of cancer invasion representing all three coculture models were used to evaluate the accuracy of invasion depth (**Figure 6A**) as well as the accuracy of binary classification whether or not the cancer cells had invaded or now (**Figure 6B**). The model had a high prediction on determining the cancer cell invasion of the three cell lines (SiHa, CaSki, and HEC-1A), showing a R^2^ of 0.9884. The binary classification model correctly identified 42 images as not invaded and 6 images as invaded. The model incorrectly identified 2 images as invaded, but it did not misclassify any invaded images. Based on these results, the model demonstrated a classification accuracy of 96%, sensitivity of 100%, and specificity of 95%.

**Figure 6.**
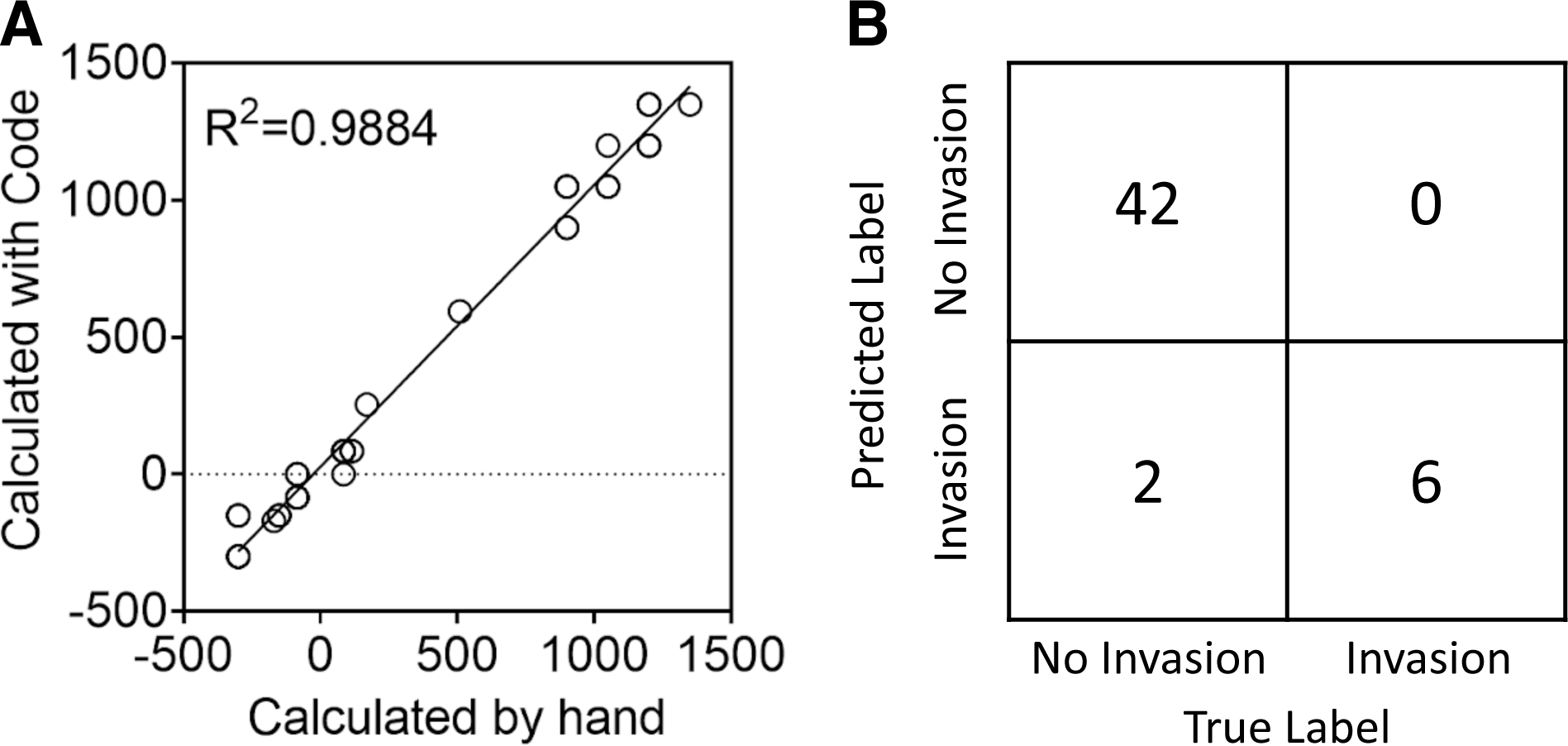
Comparison of cancer cell invasion measured by hand and with automated software. Randomly selected images representing all three coculture models were selected to assess the performance of the invasion depth code. (**A**) Comparing the invasion depth calculated with the code to the invasion depth calculated by hand. (**B**) Confusion matrix of cancer cell invasion. N=50 images.

### 3.3. Microvessel network

To assess the accuracy of our software package’s microvessel length analysis system, we compared the metrics obtained from the automated branching analysis pipeline with the metrics obtained from manual inspection (**Figure 7**). This was done using randomly selected images ranging from the minimum to the maximum of microvessel length representing all three coculture models. The R^2^ value for microvessel length exceeded 80% when comparing measurements predicted by the automated script to those obtained manually in Fiji ImageJ (**Figure 7A**). These findings are supported by the results displayed in the residual plot, where the predicted values are dispersed around 0 (**Figure 7B**).

**Figure 7.**
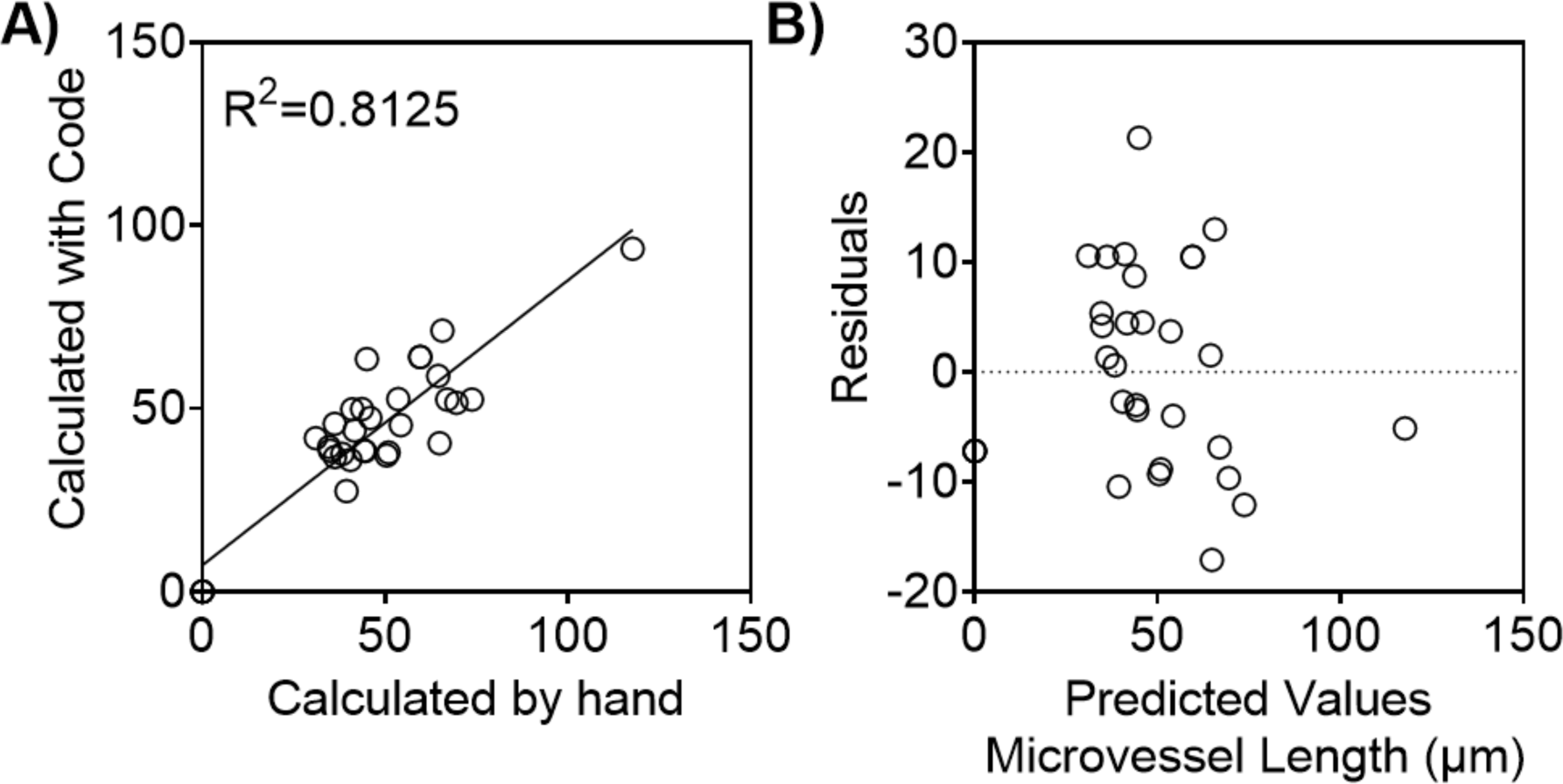
Average microvessel length measured by hand and with the automated software. Randomly selected images representing all three coculture models were selected to assess the performance of the microvessel length code. **(A)** Comparison of endothelial cell’s microvessel length measured by hand using Fiji ImageJ, and algorithm that automatically detects and quantifies the microvessel length **(B)** Residual plot showing the difference between the ground truth average microvessel length and the metrics computed by the microvessel analysis pipeline. N=32 images.

### 3.4. Comparison of dose-response analysis

To further expand the application of our automated image analysis software into drug screening studies, we performed a comparative analysis of the response of endothelial cells (hMVEC) and the cervical cancer cells (SiHa and Ca Ski) to Paclitaxel. The automated image analysis software was utilized to evaluate IC_50_ values, defined as the concentration of drug that reduced either cancer invasion, cancer coverage, microvessel length, or endothelial cell coverage to 50% of the value observed in the absence of drug. These values were then compared to the previously reported values measured manually in Fiji ImageJ and Gen5 (Cadena et al., 2023).

All of the cell lines demonstrated a similar dose-response curve compared to the previously reported values that were measured by hand (**Figure 8, Figure S3**). Consistent with our prior findings, we reported a higher IC_50_ value for cervical cancer invasion in Ca Ski cells compared to SiHa cells. Additionally, the log IC_50_ values obtained through manual measurements and our automated image analysis code demonstrated a consistent and comparable trend across the dose-response interval for Paclitaxel in both endothelial cells (hMVEC) and cervical cancer cells (SiHa and Ca Ski). These findings suggest that both approaches reliably capture the upper and lower limits of the dose-response interval (**Figure 9, Figure S4**).

**Figure 8.**
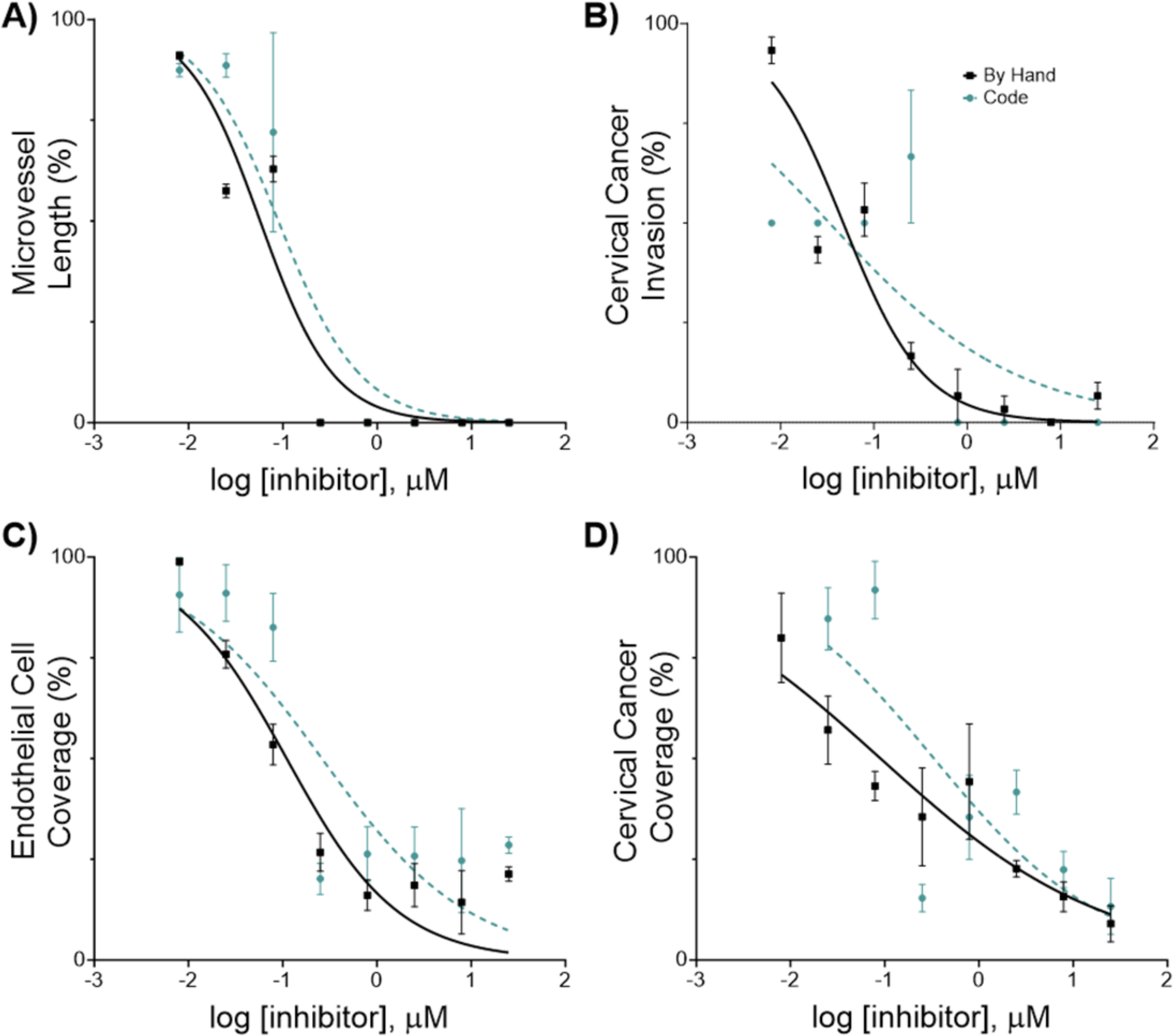
*In vitro* effects of Paclitaxel on human microvascular endothelial cells (hMVECs) and human cervical cancer cell line (SiHa). Phenotypic cell responses were measured using the automated image analysis tool (code) and using Fiji ImageJ and the Gen 5 software (by hand). Cells were co-cultured in the 3D *in vitro* model for 24 hours, then treated with 0.008 – 25 µM of Paclitaxel for 24 hours, at which point cell response was evaluated. Each cell response is normalized to the average of the values observed in the absence of the drug. **(A)** Microvessel length **(B)** Cervical cancer invasion **(C)** Endothelial cell coverage **(D)** Cervical cancer coverage. Data represent the mean ± SEM (n = 3).

**Figure 9.**
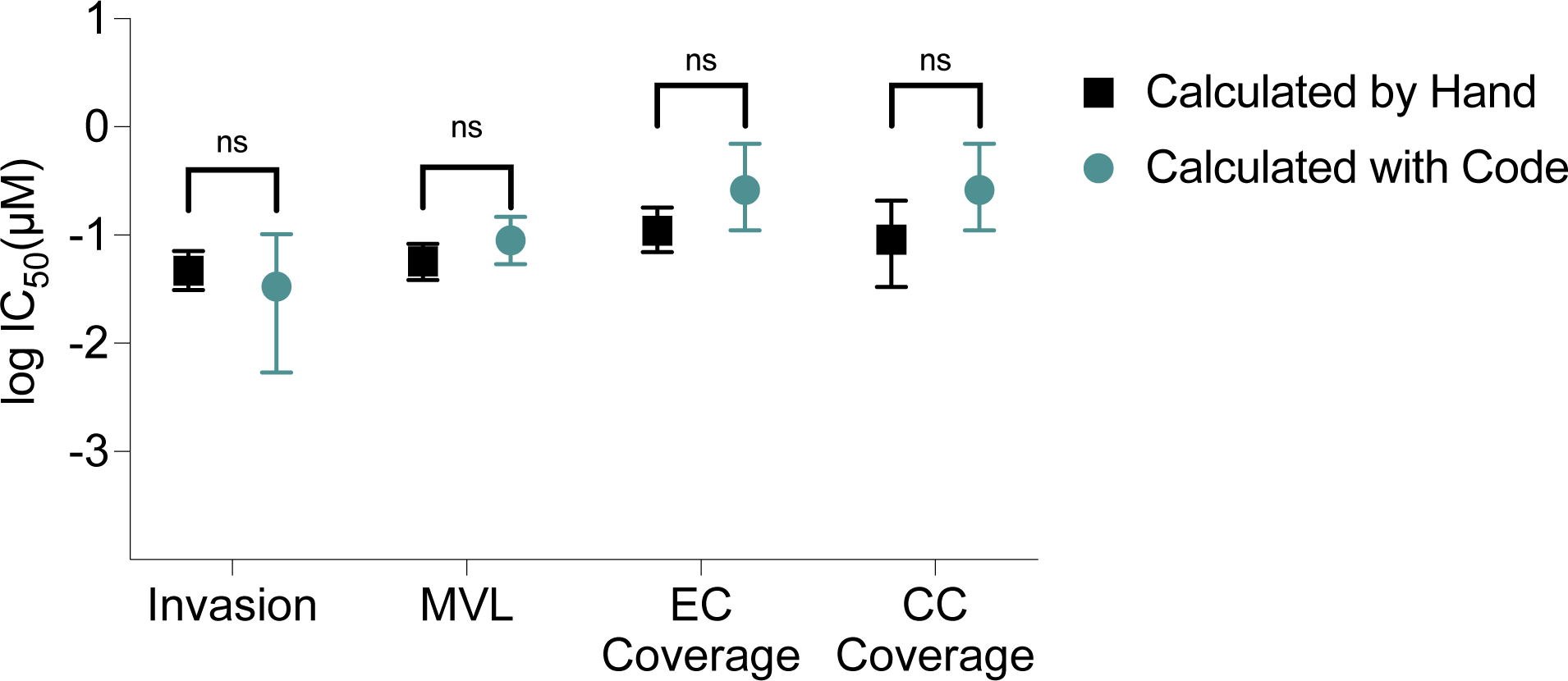
Comparison of drug inhibitory effects (IC_50_) on phenotypic cell responses. Mean values of the logIC_50_ from each inhibitory drug response curve were measured by hand and with the code in the multilayer models with endothelial cells (hMVEC) at the bottom and cervical cancer cells (SiHa) on top. Mean and upper and lower limits are represented by the error bars (N=3). Microvessel length (MVL), EC endothelial cells (EC), cervical cancer cells (CC).

## 4. Discussion

In this study, we developed an open-source Python software package designed for the automated analysis of dynamic cell behaviors within 3D tumor models. This software represents a significant advancement in high throughput screening using non-confocal images. By integrating computer vision, machine learning, and topological data analysis, we are able to quantify cell coverage area, invasion depth, and microvessel network characteristics. This suite of tools not only achieves high correlation with manual measurements but also significantly enhances the efficiency and scalability of analyzing the influence of microenvironmental cues and novel therapeutics on cancer progression.

In contrast to popular open source bioimage analysis software like Fiji ImageJ, our software is targeted to address the distinct challenges in high throughput measurement of these complex and meaningful phenotypic parameters. A key innovation in our approach lies in its ability to yield accurate results with relatively small annotated datasets, a notable departure from the norm in machine learning applications for image analysis (Nakagawa et al., 2019; Rundo et al., 2020). Through strategic data augmentation, we trained our models on fewer than 1000 binary annotations for invasion depth and 50 binary masks for microvessel segmentation. This methodology not only demonstrates the efficacy of ML in biological image analysis with limited data but also significantly reduces the manual labor involved in model training, making advanced analysis accessible to a broader range of researchers.

The capabilities and accuracy of our software package mark a major advancement in the automated analysis of 3D tumor models that are imaged in a high throughput high content manner. By achieving over 90% accuracy for cell coverage, high classification accuracy in invasion depth analysis, and greater than 95% precision in determining IC_50_ values, our tool sets a new standard for reliability in complex image analyses. This technology allows for a deeper understanding of tumor dynamics on a large scale, encompassing growth, invasion, and angiogenesis, thereby contributing to the identification and development of more effective cancer therapies. This capability not only streamlines the evaluation of drug efficacy but also substantially reduces analysis time. By potentially increasing the throughput of drug screening studies, our software paves the way for more rapid and accurate assessments of chemotherapeutic agents in 3D tumor models, enhancing the development of personalized treatment strategies.

In contrast to the “black box” nature of many end-to-end analysis tools, our software incorporates visualization throughout the process, ensuring transparency and reproducibility. Specifically, a command line flag enables saving intermediate image transformations during the cell area measurement and microvessel analysis. Experimenters can also use these graphical outputs to inform their search for the best configuration parameter values. These features not only empower users to fine-tune the software to their specific needs but also encourage engagement with the tool, fostering a collaborative ecosystem around its continued development.

Automated tools have the potential to facilitate consistent and standardized assessments of phenotypic characteristics, yet it is crucial to mitigate systematic errors within the analysis pipeline. Fine-tuning the software’s configuration parameters typically results in localized errors that can be understood and addressed through the examination of intermediate outputs. For example, parts of a vessel branch may be overlooked in sections of an image affected by noise or blur, or a minor segment of a Z-projected image might be mistakenly classified as cellular area due to structured noise artifacts inherent to the Z-projection technique. Our findings indicate that such localized inaccuracies seldom translate into significant measurement errors across an entire image, affecting neither the calculated average length of microvessel branches nor the overall cellular area substantially. Importantly, this automated approach is not prone to cognitive biases like inattentional blindness—a phenomenon in which observers overlook unexpected yet significant elements within their field of vision (Simons, 2000). This attribute of the automated system enhances the reliability of the measurements and significantly lightens the experimentalists’ burden in validating the reported phenotypic metrics.

Our image analysis software provides metrics related to cell area, invasion depth, and vessel formation, offering a glimpse into the array of phenotypic parameters available for exploration. While invaluable, these metrics capture just a fraction of the potential insights into cellular behavior. For instance, the application of persistent homology in our software facilitates the decomposition of endothelial networks into discernible branches, enabling the examination of network branching structures. Beyond mere decomposition, the mathematical framework of persistence barcodes, including well-defined distances (Cohen-Steiner et al., 2005, 2010) and kernels (Reininghaus et al., 2015) enriches the analysis, allowing for sophisticated comparisons of persistence diagrams. Such comparisons could track changes in the endothelial network’s branching structure over time or in response to treatments. The intricate mathematical structure of persistence barcodes also opens the door to applying machine learning algorithms, such as k-medoids clustering and support-vector machines, to these barcodes, suggesting exciting possibilities for future machine learning endeavors based on our software’s outputs. Moreover, our current focus on a single filtration approach for analyzing endothelial networks overlooks other valuable aspects, such as network tortuosity, highlighted by Stolz et al. (Stolz et al., 2017). Exploring alternative filtration techniques could unveil additional dimensions of network behavior, enriching our understanding of endothelial dynamics. Future developments might expand the software’s capabilities to incorporate these multifaceted analyses, further broadening the scope of phenotypic parameters that can be extracted and interpreted.

In conclusion, our open-source package represents a significant step forward in automating the analysis of 3D tumor models. By facilitating high-throughput, accurate, and reproducible analysis, it paves the way for deeper understanding and more effective interventions in cancer progression. We encourage researchers and developers to join us in enhancing and extending this tool, available at https://github.com/fogg-lab/tissue-model-analysis-tools/tree/main/, to explore the untapped potential of 3D tumor model analysis.

## Acknowledgements

We thank the Oregon State University College of Pharmacy High-Throughput Screening Services Laboratory (HTSSL). Funding was provided by Hewlett Packard (HP) to Dr. Fogg.

## Supplemental Figures

**Figure S1.**
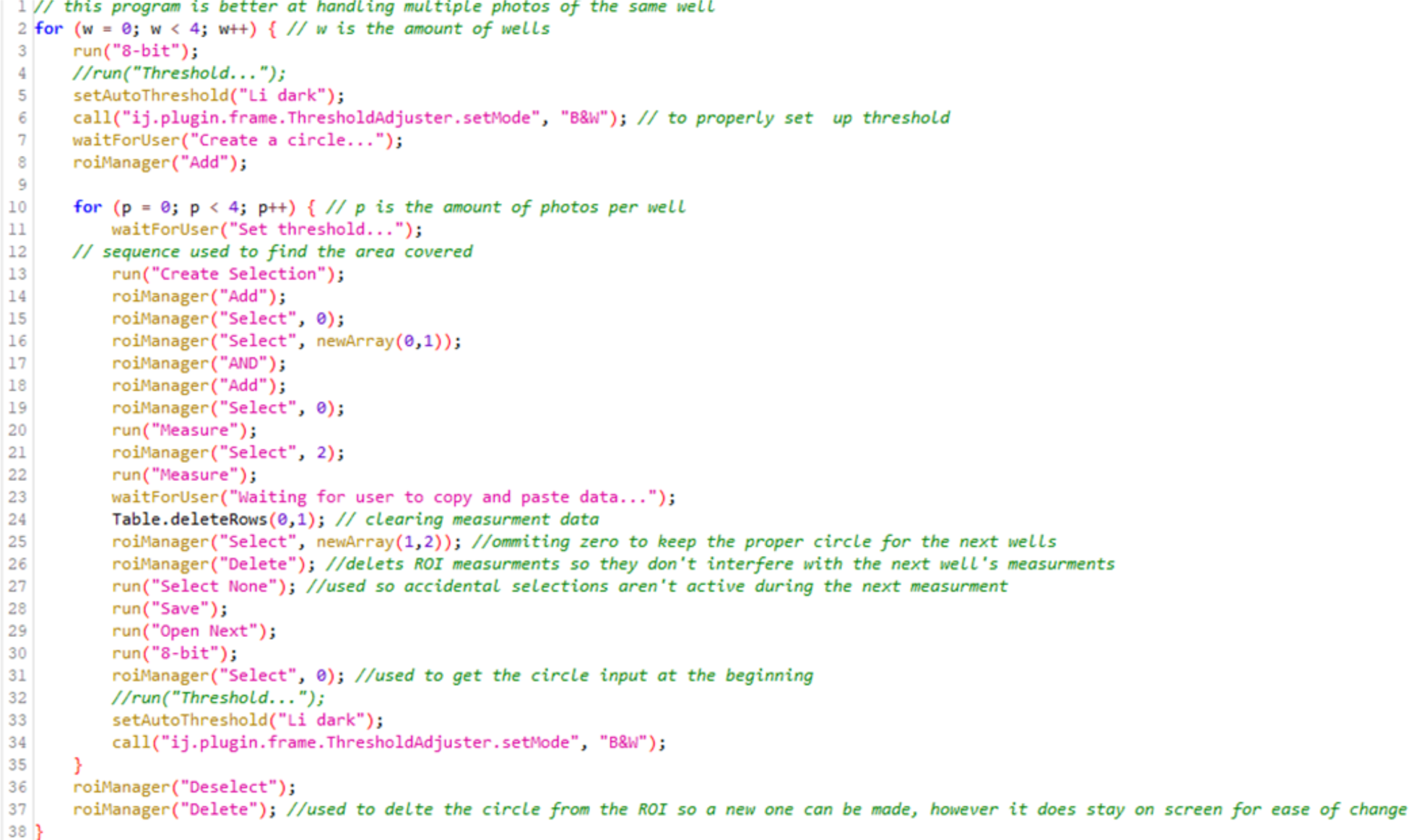
Manual inspection of cell coverage area. Using NIH Fiji-ImageJ z-projected images were processed using the Macro code to determine endothelial cell coverage and cancer cell coverage over time.

**Figure S2.**
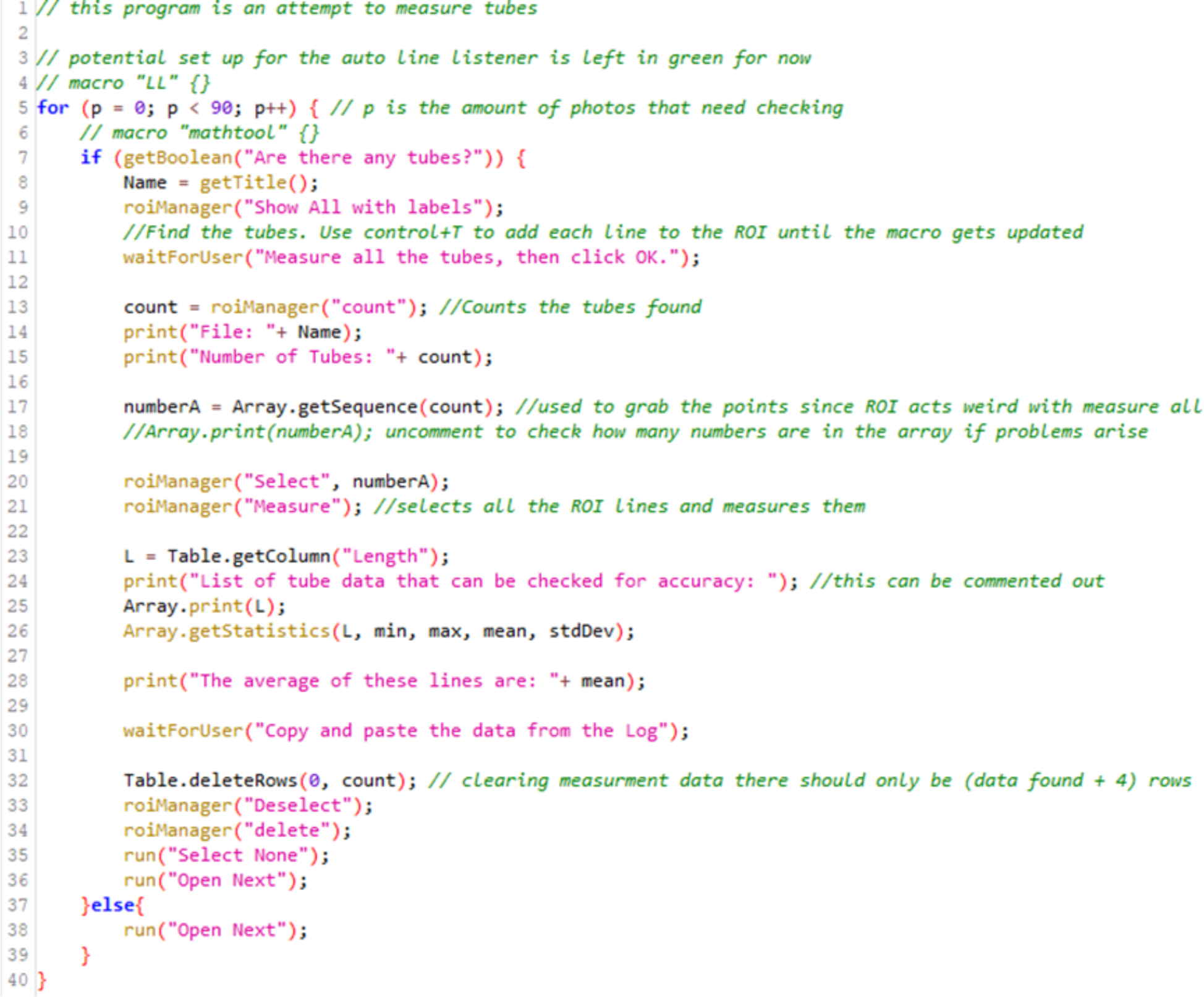
Manual quantification of microvessel length. Using NIH Fiji-ImageJ z-projected images were processed using the Macro code to determine the number and length of microvessel over time.

**Figure S3.**
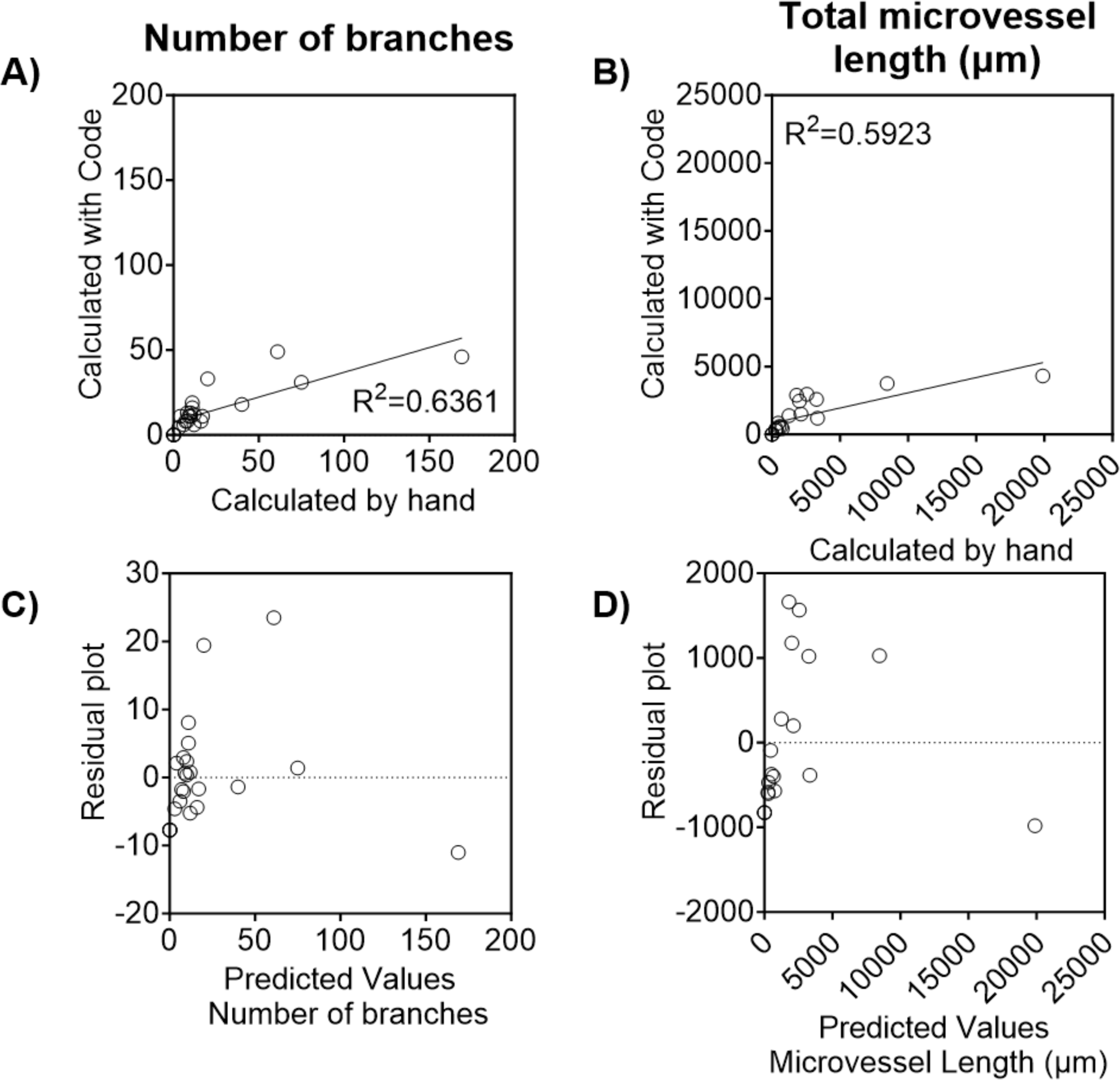
Average number of branches and total microvessel length measured by hand and with the automated image analysis software. Comparison of endothelial cell’s number of branches and microvessel length measured by hand using Fiji ImageJ, and algorithm that automatically detects and quantifies the number of branches and the microvessel length **(A, B)**. Residual plot showing the difference between the ground truth and the metrics computed by the pipeline for (**C**) number of branches, and (**D**) total microvessel length. Images were randomly selected from the 3D *in vitro* models of cervical cancer and endometrial cancer. Endothelial cells (hMVEC) were seeded on top of our optimized disease-specific hydrogels, and also on Matrigel. The different microvessel formation was quantified using the macro script developed using Fiji ImageJ and compared to the predicted values from the software with the microvessel analysis pipeline. N=32 images.

**Figure S3.**
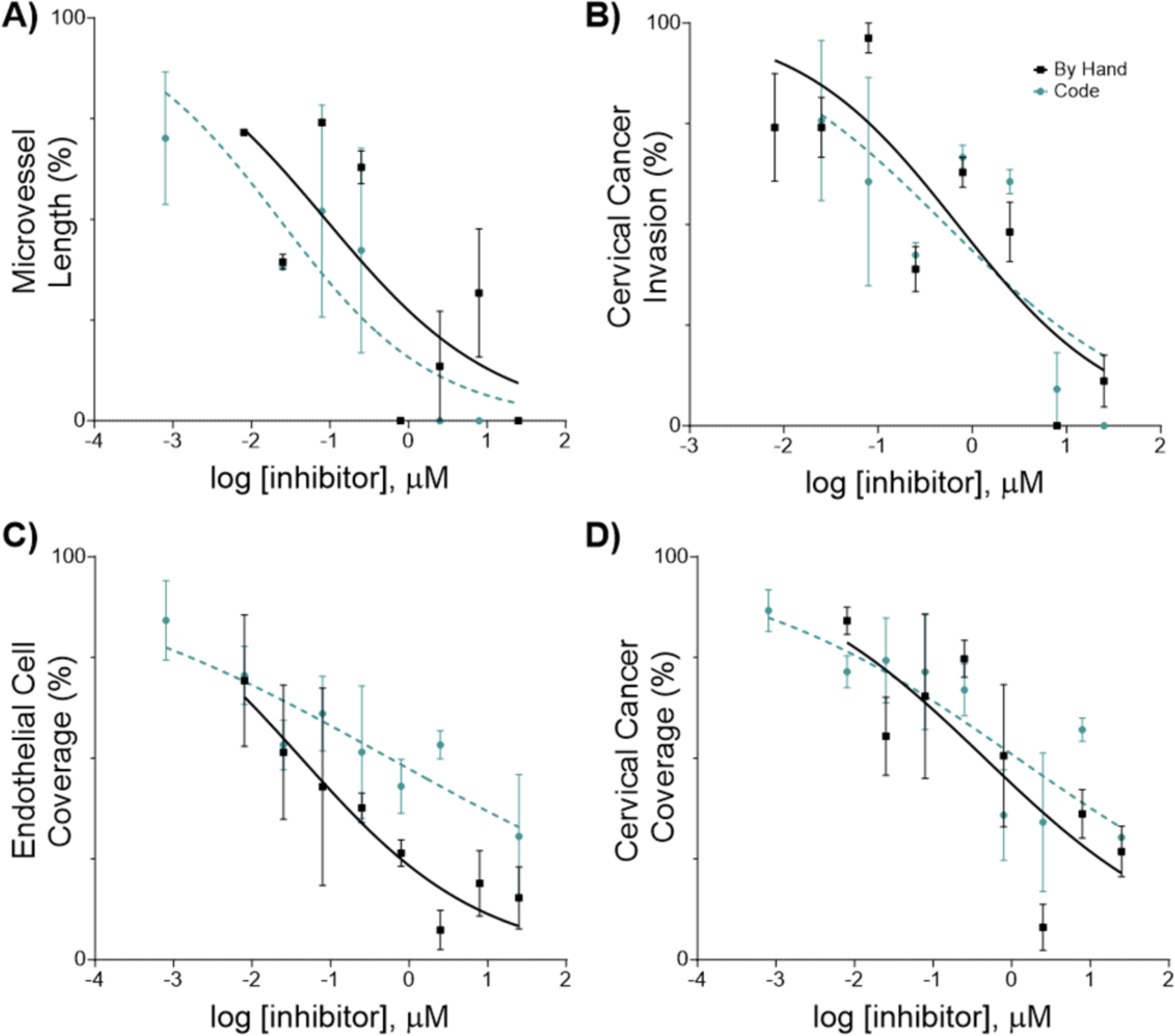
*In vitro* growth-inhibitory effects of the inhibitory drugs on human microvascular endothelial cells (hMVECs) and human cervical cancer cell line (CaSki). Phenotypic cell response measured with the code and by hand. Cells were seeded in the construct, cultured for 24 hours, then treated with 0.008 – 25 µM of Paclitaxel for 24 hours, at which point cell response was evaluated. Each cell response is normalized to the average of the values observed in the absence of drug. **(A)** Microvessel length **(B)** Cervical cancer invasion **(C)** Endothelial cell coverage **(D)** Cervical cancer coverage. Data represent the mean ± SEM (n = 3).

**Figure S4.**
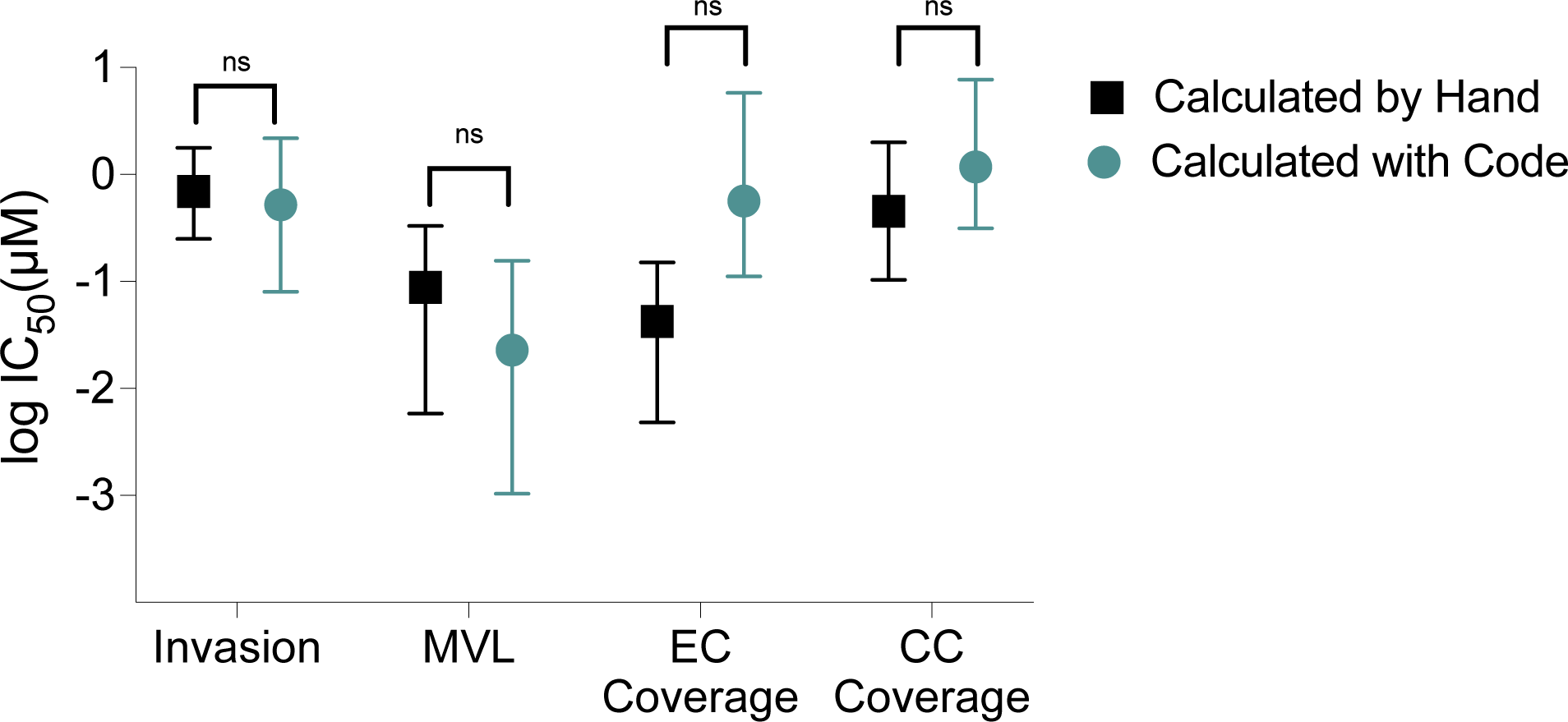
Comparison of drug inhibitory effects (IC_50_) on phenotypic cell responses. Mean values of the logIC_50_ from each inhibitory drug response curve measured by hand and with the code in the multilayer models with endothelial cells (hMVEC) in the bottom and cervical cancer cells (Ca Ski) on top. Mean and upper and lower limits represented by the error bars (n=3). Microvessel length (MVL), endothelial cells (EC), cervical cancer cells (CC).

